# The importance of habitat type and historical fire regimes in arthropod community response following large-scale wildfires

**DOI:** 10.1101/2023.07.17.548903

**Authors:** Anna J. Holmquist, R.J. Cody Markelz, Ciera C. Martinez, Rosemary G. Gillespie

**Author notes:** Correspondence: Anna J. Holmquist, Center for Comparative Genomics, California Academy of Sciences, San Francisco CA USA.

## Abstract

Wildfires are increasingly altering ecosystems, posing significant challenges for biodiversity conservation and ecosystem management. In this study, we used DNA metabarcoding to assess the response of arthropod communities to large-scale wildfires across diverse habitat types. We sampled six reserves within the University of California Natural Reserve System (UCNRS), each which was partially burned in the 2020 Lightning Complex wildfires in California. Using yellow pan traps to target pollinators, we collected arthropods from burned and unburned sites across multiple habitat types including oak woodland, redwood, scrub, chamise, grassland, forest, and serpentine habitats. We found no significant difference in alpha diversity values between burned and unburned sites; instead, seasonal variations played a significant role in arthropod community dynamics, with the emergence of plant species in Spring promoting increased pollinator richness at all sites. Compositional similarity analysis revealed that burn status was not a significant grouping factor when comparing all sites. Instead, community composition primarily varied across reserves, indicating distinct pools of arthropods structured geographically. Habitat type played a crucial role in determining the response of arthropod communities to fire. While communities in grasslands and oak woodlands exhibited recovery following burn, scrublands experienced substantial changes in community composition. Our study highlights the importance of examining community responses to wildfires across broad spatial scales and diverse habitat types. By understanding the nuanced dynamics of arthropod communities in response to fire disturbances, we can develop effective conservation strategies that promote resilience and maintain biodiversity in the face of increasing wildfire frequency and intensity driven by climate change.

## Introduction

Disturbance is integral to the health and productivity of many ecosystems. The intermediate disturbance hypothesis (IDH) encapsulates this, with the prediction that diversity will be highest under intermediate levels of disturbance (Connell 1978; Roxburgh, Shea, and Wilson, J.B. 2004). However, with climate change comes an increasing number of severe and devastating natural disasters that are shifting established disturbance regimes and leading to rapid shifts in ecosystems (Johnstone et al. 2016; Seidl et al. 2017). Wildfires are one such example; while many ecosystems are adapted to cyclical fire regimes that have been maintained over millions of years (Pausas and Keeley 2019; 2009), specific aspects of wildfire regimes are changing due to shifts in climate as well as increases in anthropogenic ignition (Jager et al. 2021; Eds 2007). In the Western USA, both fire frequency, area, and severity have seen sharp increases since the 1980s (Parks and Abatzoglou 2020). With fire activity predicted to continue intensifying alongside climate change, effective management requires that we understand how the regime shifts are impacting biotic communities (Schoennagel et al. 2017).

Many habitat types and their associated biota rely on fire to maintain an ecological steady state. Historical regimes generally featured low severity fires at consistent intervals that generated a heterogeneous landscape; the resulting variability in resources and vegetation structure can create new niche space that increases species diversity (Taillie et al. 2018). Natural wildfire regimes produce a range of supporting, provisioning, regulating, and cultural ecosystem services for humans (Pausas and Keeley 2019). However, modern regimes featuring frequent and high-intensity fires are driving forest conversion even in habitat types accustomed to fire such as in conifer-dominated forests (Coop et al. 2020) and pine-oak woodlands (Barton 2002). Intense wildfires can not only kill large expanses of trees but additionally alter soil quality, reduce seed reserves, and facilitate the establishment of non-native plant species. Such catastrophic wildfires can change ecosystem structure and cause the loss of important ecosystem services such as erosion control and carbon sequestration (Lecina-Diaz et al. 2021).

The effects of fire on local fauna are also influenced by its severity. Plant-pollinator diversity has been shown to benefit from natural wildfire, with the highest richness of plants and insect pollinators found in sites that experienced low to moderate severity wildfires; however, this trend is lost in areas of high-severity burn (Ponisio et al. 2016). A recent meta-analysis found a strong effect of high-severity fires on arthropod communities as a whole, consistently resulting in a reduction in richness and evenness (Bieber et al. 2022). Another meta-analysis found increases in pollinator abundance and richness after fire, but found shortened fire intervals to be detrimental to communities (Carbone et al. 2019). Both papers emphasize the importance of dynamics of the fire regime, such as fire intensity and burn interval, rather than burn itself, on the arthropod community. Habitat type (e.g. grasslands, desert, forest) has been identified as an additionally important factor in dictating the response of arthropod communities to fire (Bieber et al. 2022) although most studies thus far have focused on forested areas. The response of vegetation and the rest of the biotic community is largely dependent on habitat type and historical fire adaptation. While habitats adapted to fire can still be severely impacted by high-intensity fires, habitat types that rarely experience fire are at higher risk of reaching ecosystem tipping points following a burn (Lecina-Diaz et al. 2021; Moritz et al. 2012). Understanding broad patterns of community response following wildfire therefore requires assessment across diverse habitat types.

Because of the tight association between many insect species and the floral community as well as their importance in bottom-up structuring of the trophic network, insects can serve as excellent bioindicators of local ecosystem state following disturbance. Moreover, insects provide an opportunity to assess whole-community response through their ease of capture, providing a more generalizable picture of the effects of disturbance than single-taxon studies. DNA metabarcoding approaches have enabled large-scale studies of arthropod communities and how community structure changes following different types of disturbance, including establishment of invasive taxa (Holmquist, Adams, and Gillespie 2023), land-use changes (Beng et al. 2016) and urbanization (Dürrbaum et al. n.d.). Our study aims to address the response of arthropod communities to another major disturbance – wildfires – across diverse habitat types that differ in historical exposure to fire. We focused on protected natural areas that are part of the University of California Natural Reserve System (UCNRS), many of which were at least partially burned during the 2020 August Lightning Complex Fire. This fire system caused over 650 wildfires across the state, and burned 1,500,000 – 2,100,000 acres of land (CAL Fire 2022). It is one of many recent large-scale fires that have set records for California (CAL Fire 2022). The UC reserves, distributed across most of the major California ecoregions, offer a unique opportunity to examine fire effects on arthropods across a broad spatial scale in distinct habitat types that differ in historical fire regimes. The seven habitat types in our study are grassland, scrub, oak woodland, forest, redwood, chamise and serpentine. Each of these habitat types has a different pre-settlement fire interval estimate in years: grassland (< 10), oak woodland (12), redwood (23), mixed pine forest (23), and scrub (60) (Pausas and Keeley 2019). This allowed us to test how differences in historical fire frequency in the different habitat types can alter community response.

To measure the response of arthropod communities to fires, we adopted a DNA metabarcoding approach to sequence all individuals in a given community. Metabarcoding techniques allow rapid classification of biotic communities without the need for taxonomic identifications. Adoption of a non-destructive extraction approach additionally allows *post-hoc* assessment of abundances and verification of results. We targeted pollinator communities specifically because of their importance in plant diversity as well as their reliance on floral communities. To do this, we used yellow pan traps to sample across burned and unburned sites in Fall 2020 and Spring 2021 following the major Lightning Complex wildfires in California. We used these data to test a) how species richness varies between burned and unburned sites, b) the extent of compositional differences between burned and unburned sites and c) the role of habitats with different historical fire regimes in shaping community response. Our specific hypotheses were:

1. Because many insect species are adapted to fire and will colonize following burn, alpha diversity will not differ between burned and unburned sites. Differences in alpha diversity will instead be associated with season due to changes in plant activity and the associated emergence of pollinators.
2. Community recovery from fire depends on re-colonization, which is in part driven by dispersal capabilities. Because we are targeting pollinators, most of which are winged taxa and dispersive, we expect no strong compositional differences when comparing sites within the same habitat types between different reserves. Compositional differences between sites will be driven instead by season, habitat type, and/or burn status.
3. The extent of dissimilarity between burned and unburned sites will differ by habitat type, with habitat that rapidly recovers from any major disturbance (grasslands) or habitat well-adapted to fire (oak woodlands) showing recovery by the Spring, while habitat less accustomed to severe wildfire (scrub) will remain altered compared to unburned sites.

## Methods

### UC Reserves

The UC Natural Reserve System comprises 47,000 acres distributed across most of the major California ecosystems. Between August 15-16, 2020, an exceptionally fierce electrical storm crossed northern California, peaking at 200 lightning strikes in a 30-minute period. Over the following 72–96 hours, more than 12,000 lightning strikes sparked multiple wildfires around the Greater Bay Area. Many of the smaller fires merged into four large fire complexes, including two in Monterey County. Three of these fire complexes are considered to be the second, third, and fourth largest fires in California history (CAL Fire 2022)^20^. These fires burned eight of the UC Natural Reserve System’s 41 reserves, and consumed more than 20,000 acres of reserve lands. Affected NRS reserves include: Año Nuevo Island Reserve (CZU Lightning Complex), Blue Oak Ranch Reserve (SCU Lightning Complex), Landels-Hill Big Creek Reserve (Dolan Fire), Hastings Natural History Reservation (River Fire), McLaughlin Natural Reserve (LNU Lightning Complex), Point Reyes Field Station (Woodward Fire), Quail Ridge Reserve (LNU Lightning Complex), and Stebbins Cold Canyon Reserve (LNU Lightning Complex). A UC Davis property affiliated with Stebbins Cold Canyon, the Cahill Reserve, also burned extensively due to the LNU Lightning Complex. The current study sampled six burned reserves: Quail Ridge, McLaughlin, Hastings, Blue Oak, Big Creek, and Ano Nuevo.

### Field collection

A joint effort between the University of California: Berkeley and the University of California: Santa Cruz was undertaken to sample plant and arthropod communities across the six burned reserves (Figure 1). At each reserve, burned and unburned sites were selected within eight habitat types: oak woodland, redwood, scrub, chamise, grassland, forest, and serpentine. Arthropods were collected at each site using yellow pan traps filled with water and added soap to break surface tension. Yellow pan traps typically attract pollinators, but can attract multiple other insect types; all taxa were retrieved from pan traps. Fall 2020 pan traps were placed in October – December and Spring 2021 pan traps were placed in May – June. Traps were deployed for one day and moved to whirlpaks following the sampling period. Samples were sorted and stored in ethanol at -20°C until further processing.

**Figure 1.**
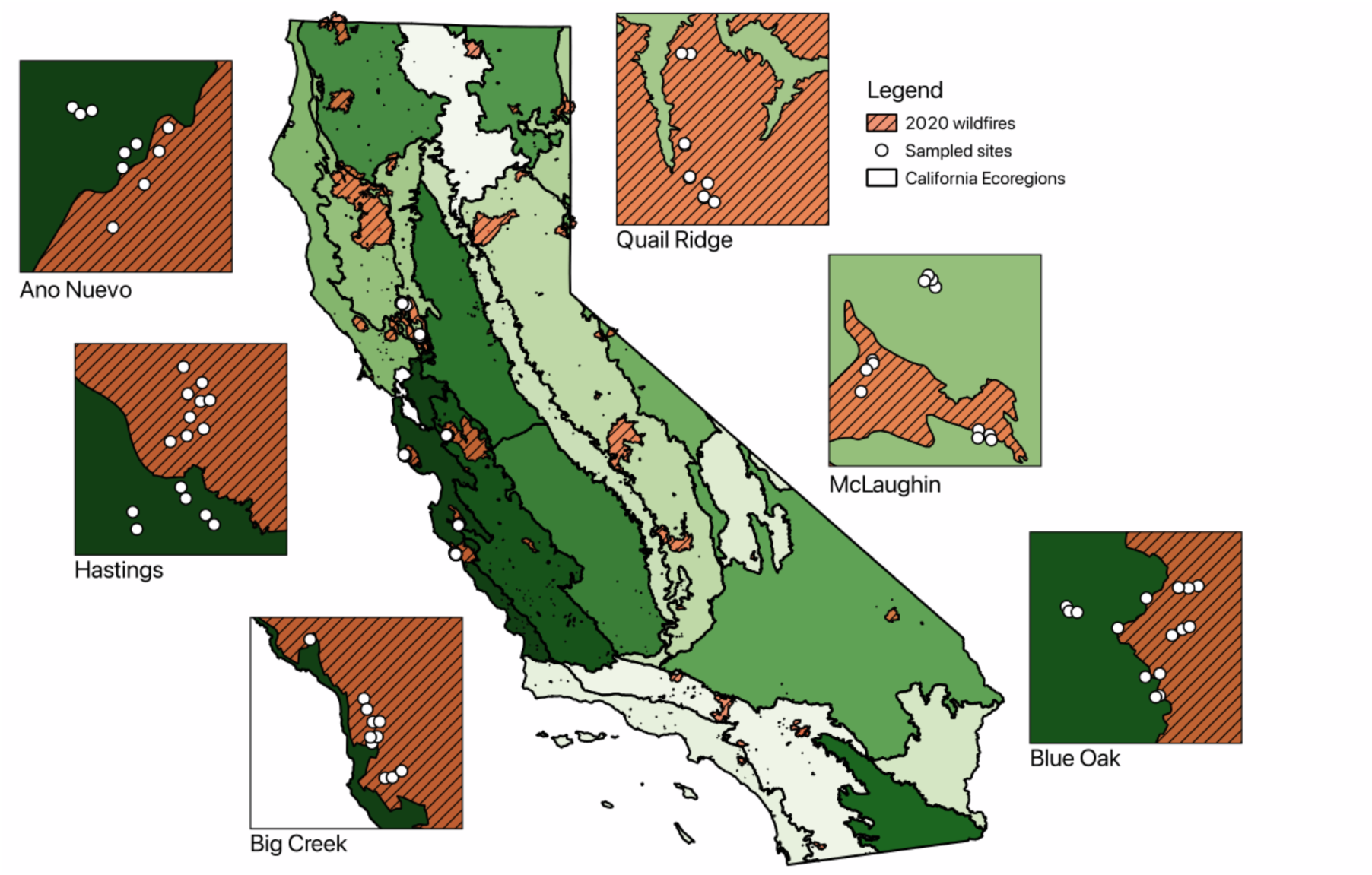
Map showing the California Lightening Complex fires of 2020, as well as points (red stars) highlighting the reserves sampled in our study. Reserves shown are: Quail Ridge, McLaughlin, Hastings, Blue Oak, Big Creek, and Ano Nuevo.

### Molecular procedures

All arthropods collected from pan traps were used in this study. Samples were counted then subdivided into hard-bodied and soft-bodied arthropods to allow varied lysing time. Once sorted, samples were stored in 95% EtOH in 96-well plates at -20°C. A BenchSmart 96 (Mettler-Toledo Rainin, Colombus, OH, USA) was used to remove EtOH after spinning plates down briefly. Remaining EtOH was evaporated using a vacuum centrifuge, and checked at regular intervals until fully dry. Extraction protocols were modified from the Qiagen PureGene extraction kit (Qiagen, Hilden, Germany). Hard-bodied insects were incubated for 24 hours at 55°C in cell lysis buffer and Proteinase K while soft-bodied insects were incubated for 3 hours. Following incubation, lysate was transferred to a second plate and RNAse added, followed by an additional incubation period at 37°C for 30 minutes. Plates were then placed on ice and, once cool, Protein Precipitation Solution was added. Plates were spun for 10 minutes at 2000rcf to form protein pellets, then supernatant was transferred to a solution of isopropanol and glycogen to precipitate DNA. Plates containing specimens were washed repeatedly to remove lysate and stored in 70% EtOH for future taxonomic work. Samples were incubated at room temperature overnight then spun down to form DNA pellets. Isopropanol was removed and pellets were washed with cold 70% EtOH. Ethanol was removed and, once remaining ethanol evaporated, elution buffer was added and then plates incubated at 65°C for 1 hour. Plates were then aliquoted and stored at – 20°C until further use.

Amplification of COI was performed using the Qiagen multiplex kit (Qiagen, Hilden, Germany). Two sets of primers were used – MCO and BF3 (Table 1) (Leray et al. 2013; Elbrecht et al. 2019; Yu et al. 2012). Primers additionally had a 5#x2019; TruSeq tail for indexing. Amplification was performed on 96-well plates and consisted of 5ul of the Qiagen PCR MasterMix (MM) (Qiagen, Hilden, Germany), 3ul of H_2_0, 0.5ul of each primer, and 1ul of template DNA. Three PCR replicates were performed. Annealing temperature was 46°C and ran for 30 cycles. A negative PCR control was included on each plate, consisting of MM, H20, and primers. PCR products were visualized using gel electrophoresis on a 3% agarose gel. A dual indexing strategy was implemented using a second round of PCR to attach 8bp indexes. Annealing temperature for indexing PCR was 55°C and ran for 6 cycles. PCR products were visualized once again using a 3% agarose gel and ran at low voltage to allow clear visualization of length to confirm the addition of indexes to each amplicon. Final libraries were constructed by pooling PCR products proportionally based upon band strength. Further processing was conducted at Berkeley QB3 Genomics (QB3, Berkeley, CA USA). Libraries were sequenced using Illumina MiSeq paired-end sequencing and demultiplex at QB3.

**Table 1.**
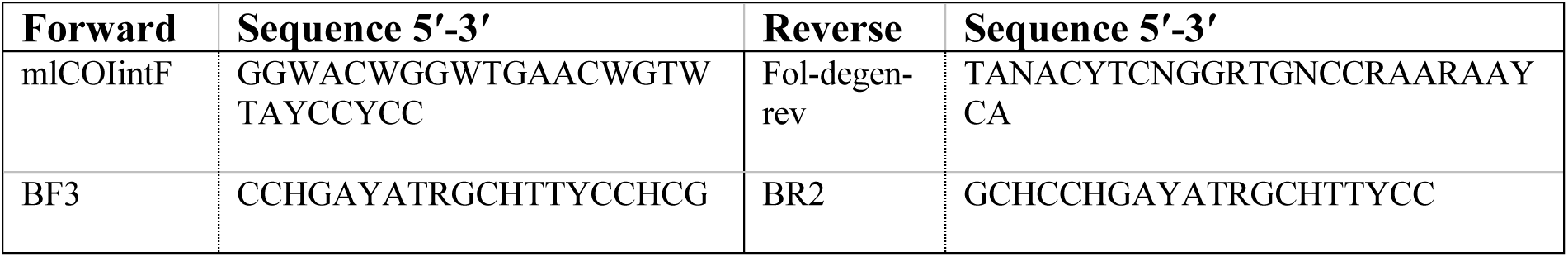
Primer pairs used to amplify COI.

### Bioinformatic processing

Reads were trimmed of primer adapters using CutAdapt (Martin 2011). Reads were processed using the DADA2 denoising algorithm(Callahan et al. 2016) to form amplicon sequence variants (ASVs). The *removeBimeraDenovo* function included in the DADA2 package was used to remove chimeras. The BF3 primer pair produced low quality sequences with over half being discarded through normal filtering procedures, and was not included in further analysis. The PCR negative controls were utilized for decontamination, using the package decontam (Davis et al. 2018). Following decontamination, the curation algorithm LULU (Frøslev et al. 2017) was used to remove erroneous sequences. Sequences were aligned in Geneious Primer Prime v.2022.0.2 to detect any remaining pseudogenes, identified by stop codons or shifts in the reading frame. Following removal, BLAST was performed using megablast. A custom script in R using the package rentrez (Winter 2017) was then used to generate full lineage information from accession numbers. Only phylum Arthropoda was retained, and sequences with below 80% percent identity matches were also removed. The final filtering step retained only ASVs found in two of three replicates. Operational taxonomic units (OTUs) were generated using *otu* in the package kmer (Wilkinson, S 2018), clustered at 97% with kmer = 5. This cluster threshold was used to examine taxa at a coarser level, closer to genus than species.

### Statistical analysis

The BF3/BR2 primer pair was found to generate high errors with over half of reads associated with the primer set being discarded when using DADA2. Because of this, only amplicons produced from the MCO primer pair were used. Reads were summed for each ASV detected across replicates for a particular sample and a community matrix was created. Because reads do not translate directly to abundance count, read data was transformed using Hellinger standardization; this method has been widely adopted in metabarcoding. Standardization was performed using *decostand* in vegan (Oksanen et al. 2009)^32^.

Quail Ridge had no unburned sites and was excluded from the analysis. The number of sites in redwood (2 burned and 1 unburned), chamise (3 burned and 2 unburned), forest (2 burned and 2 unburned), and serpentine (2 burned and 2 unburned) was low; scrub, grassland and oak woodland habitat types were the best sampled, so the data were narrowed to include only these habitat types.

We used OTU richness, the Shannon index and the Simpson index to calculate alpha diversity metrics, with the standardized read matrix as input. F-tests were performed to compare variances; the non-parametric Mann-Whitney U test was used to assess median differences between alpha diversity of burned and unburned sites within the same reserve, habitat type, and season. For compositional comparisons, only sites with ≥ 3 OTUs were used. To isolate the effect of wildfire on community differences, comparisons were made between unburned and burned sites found within the same reserve, habitat type and within the same season (Fall 2020 or Spring 2021). All analyses were performed using both incidence and Hellinger-standardized read data. Nonmetric multidimensional scaling was performed using Hellinger distance, by applying Euclidean distance to the Hellinger-standardized data; this was done using *metaMDS* in vegan with up to 1,000 random starts and 3 dimensions. The argument -noshare was used which creates extended dissimilarities for sites that share no species. Outlier samples with entirely distinct communities prevented convergence (Supplementary Figure 1) and were removed from NMDS, but all statistical tests were run using the data with and without the outlier data. Permutational multivariate analysis of variation (PERMANOVA) was used to test group differences based on burn, habitat type, reserve and year using the Hellinger distance. This was done using *adonis2* in vegan (Oksanen et al. 2009). Differences in group dispersions were tested using PERMDISP2 with the function *betadisper*. Additionally, we split the data by habitat type, and performed NMDS, PERMANOVA and PERMDISP2 for each individual habitat type.

To calculate compositional differences in an additional way, we used *beta* in the package BAT (Cardoso, Rigal, and Carvalho 2015), which partitions beta diversity into differences driven by species replacement and by richness differences. We performed the non-parametric Kruskal-Wallis test to assess if burned and unburned sites had higher beta diversity values than sites within burned habitat and sites within unburned habitat. We compared differences in beta diversity against distances between sites within reserves to examine spatial autocorrelation and the key components driving dissimilarities. Spatial distance between sites was calculated using QGIS v.3.22.4 and the standard setting under distance matrix calculation. We performed linear regression to assess the association with distance. Using ANOVA, we tested the difference between beta diversity values across habitat types.

The various comparisons used in the analyses are summarized in Table 2. Data processing and statistical analysis was performed in the R programming environment (R Core Team, 2022). Figures were created using packages ggplot, and venn. The map (Figure 1) was developed using data produced by CAL Fire.

**Table 2.** Summary of analysis and the comparisons made.

| Metric | Comparison |
| --- | --- |
| *Data summary:*<br>Taxonomy | All sites |
| *Hypothesis 1:*<br>OTU Richness, Shannon,<br>Simpson | All sites from scrub, oak woodland and grassland habitat types |
| *Hypothesis 2:*<br>NMDS, PERMANOVA,<br>PERMDISP2 | Sites with $\geq 3$ OTUs from scrub, oak woodland and grassland habitat type |
| *Hypothesis 2:*<br>Beta diversity | Sites with $\geq 3$ OTUs within the same habitat type, reserve and season |
| *Hypothesis 3:*<br>NMDS, PERMANOVA,<br>PERMDISP2 by habitat | Create separate data sets for scrub, oak woodland and grasslands habitat types; compare sites with $\geq 3$ OTUs |
| *Hypothesis 3:*<br>Beta diversity by habitat | Create separate data sets for scrub, oak woodland and grasslands habitat type; compare sites with $\geq 3$ OTUs within the same reserve and season |

## Results

### Data summary

A total of 6,461,595 reads were produced after primer removal and initial filtering by CutAdapt. Filtering, clustering and chimeral removal with DADA2 produced 5,215,076 reads. After implementing LULU, removing potential contaminants using decontam, removing NuMts, selecting only phylum Arthropoda, and removing ASVs not found in two of the three replicates for a sample, the dataset consisted of 2,167,382 reads and 584 ASVs that clustered into 448 OTUs. Taxonomic identities were assigned based on percent identity; 12 orders (> 85%), 33 families (> 92%), 94 genera (> 97%) and 51 species (>99%) were identified using BLAST.

The most common order by OTU in the Fall was Diptera and in the Spring was Hymenoptera (Figure 2). 17 families were shared across burned and unburned sites. There were nine unique families detected in unburned sites and six unique families detected in burned sites when comparing all sites. Two arachnid families were detected in burned sites (Acariformes: Erythraeidae and Araneae: Lycosidae). In addition, two unique families of dipterans (Chironomidae and Syrphidae), one unique family of lepidopterans (Lycaenidae) and one unique family of hymenopterans (Megaspilidae) were identified in burned sites. Four families of flies, two families of wasps, one family of butterflies, one family of beetles and one family of spiders were the distinct families in unburned habitat. However, only 133 of 584 OTUs were confidently identified to family with >95% identity matches.

**Figure 2.**
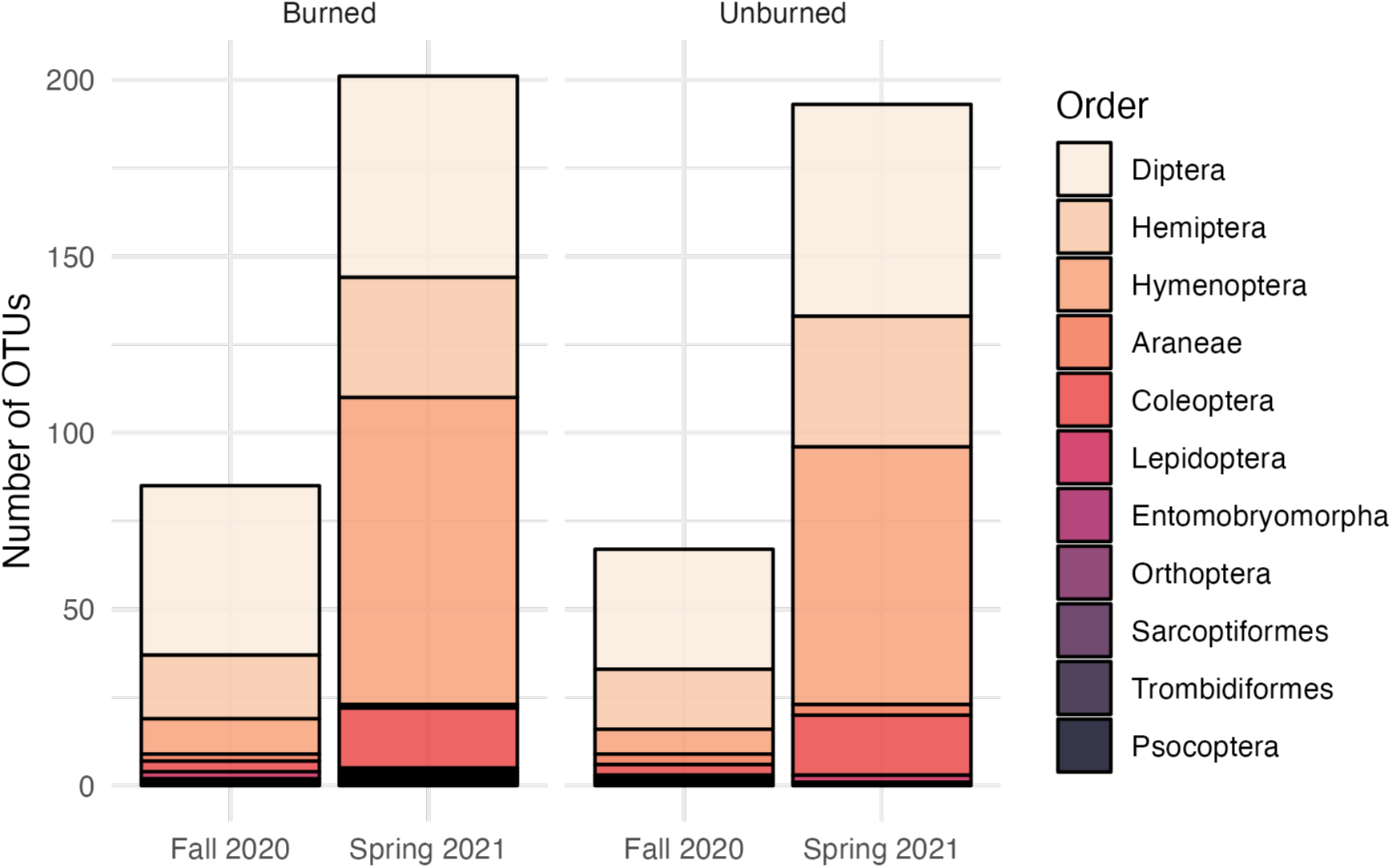
Bar diagram showing the number of OTUs belonging to each order in Fall 2020 and Spring 2021 for all burned and unburned sites. Composition was similar across burned and unburned sites. There were major increases in Hymenoptera OTUs in the spring.

### H1. Alpha diversity

Variances between burned and unburned sites were not significantly different for any of the calculated alpha diversity values. Values of alpha diversity were not significantly different between burned and unburned sites (p-value = 0.315 richness; 0.252 Shannon; 0.217 Simpson). There were significant differences by season, with an increase in richness in the Spring (p-value = 0.0006, richness; 0.0004, Shannon; 0.0009, Simpson). All habitat types saw increases in richness in the Spring compared to Fall (Figure 3).

**Figure 3.**
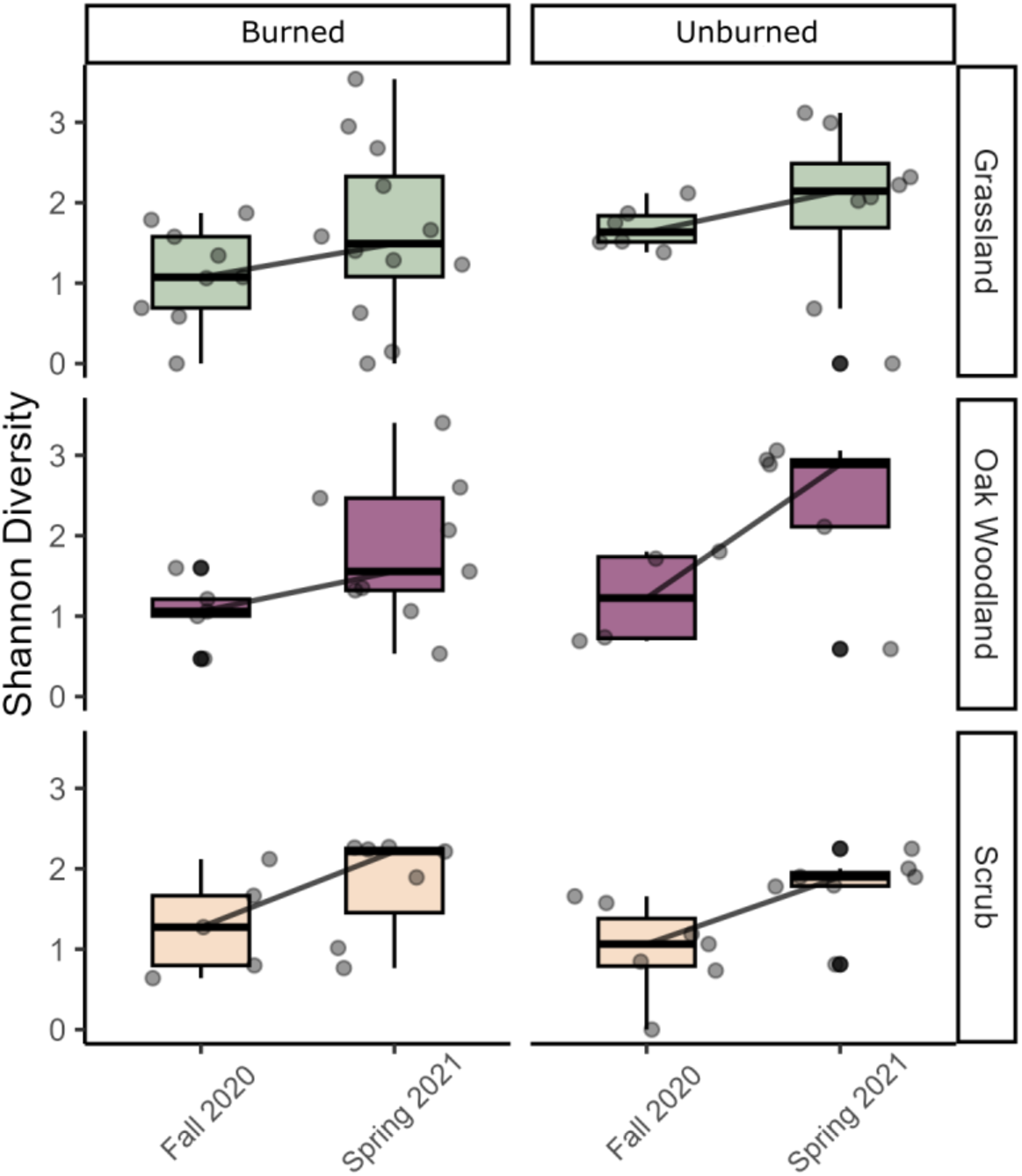
Differences in richness across fall and spring for three different habitat types (grassland top, oak woodland middle and scrub bottom) in burned sites (left) and unburned sites (right).

### H2. Predictors of community compositional differences

Results were similar between Hellinger-standardized read data and Jaccard distances (Supplementary Table 1). Because of this, results presented are from Hellinger-standardized data. Two outlier samples were removed to allow convergence of nMDS; including the outliers from statistical tests did not change the results (Supplementary Table 1, 2).

There were no significant dispersion differences between burned and unburned sites (F = 0.355, Pr = 0.554; Figure 4). Multiple variables (season, habitat type, reserve, and burn status) as well as interactions between variables were significant predictors of compositional differences between sites. Reserve was the strongest explanatory variable of group variation (R2 = 0.0925, F = 1.98, Pr < 0.001) followed by an interaction between reserve and season (R2 = 0.0799, F = 1.71, Pr < 0.001). There was a significant interaction between burn status and reserve as well (R2 = 0.0587, F = 1.26, Pr < 0.001) followed by an interaction between habitat and reserve (R2 = 0.0572, F = 1.22, Pr < 0.001). Season individually explained less variation in the response (R2 = 0.034) but was the strongest predictor of group differences based on the F-statistic (F = 2.895, Pr < 0.001; Figure 4). Burn status was not a significant explanatory variable by itself, and only showed significance when included in interactions.

**Figure 4.**
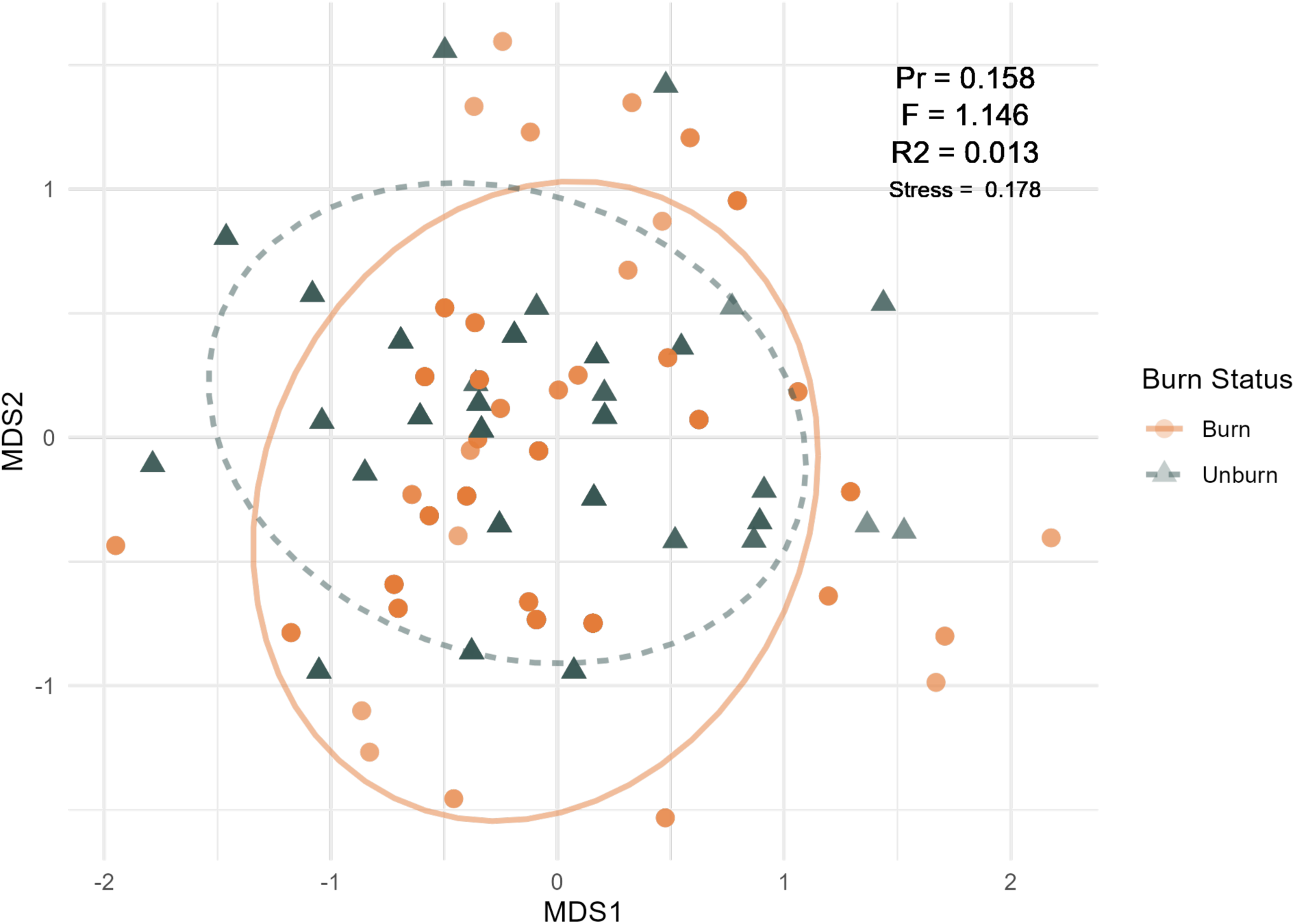
NMDS plot showing similarity of arthropod communities between sites, colored by burn status with burned sites in orange and unburned sites in green. There was no significant difference by burn status.

The median beta diversity value was 0.964 between burned and unburned sites found within the same reserve, habitat type, and during the same season. When comparing sites within unburned habitat, the median beta diversity score was 0.908 and when comparing sites within burned habitat, the median beta diversity score was 0.924. The differences between comparisons, however, were not significant. The major component of beta diversity was species replacement with a mean of 0.769 of the total beta diversity values. In the Fall, there was a slightly significant decrease in beta diversity across distance (Adjusted R2 = 0.140, t = -2.285, Pr = 0.031), driven by replacement (Adjusted R2 = 0.133, t = -2.236, Pr = 0.035) (Figure 6). This trend was lost in the Spring; instead, there was a slightly significant association with the richness component of beta diversity across distance (Adjusted R2 = 0.077, t = -2.024, Pr = 0.051), in which sites closer to one another have compositional differences driven more by differences in richness rather than differences in species present. The proportion of beta diversity explained by richness decreased over distance, with the farther sites having beta diversity values largely due to species replacement. There is a decrease in replacement over distance, although this result was not significant likely due to a few intermediate-distance sites with a richness component (Figure 5).

**Figure 5.**
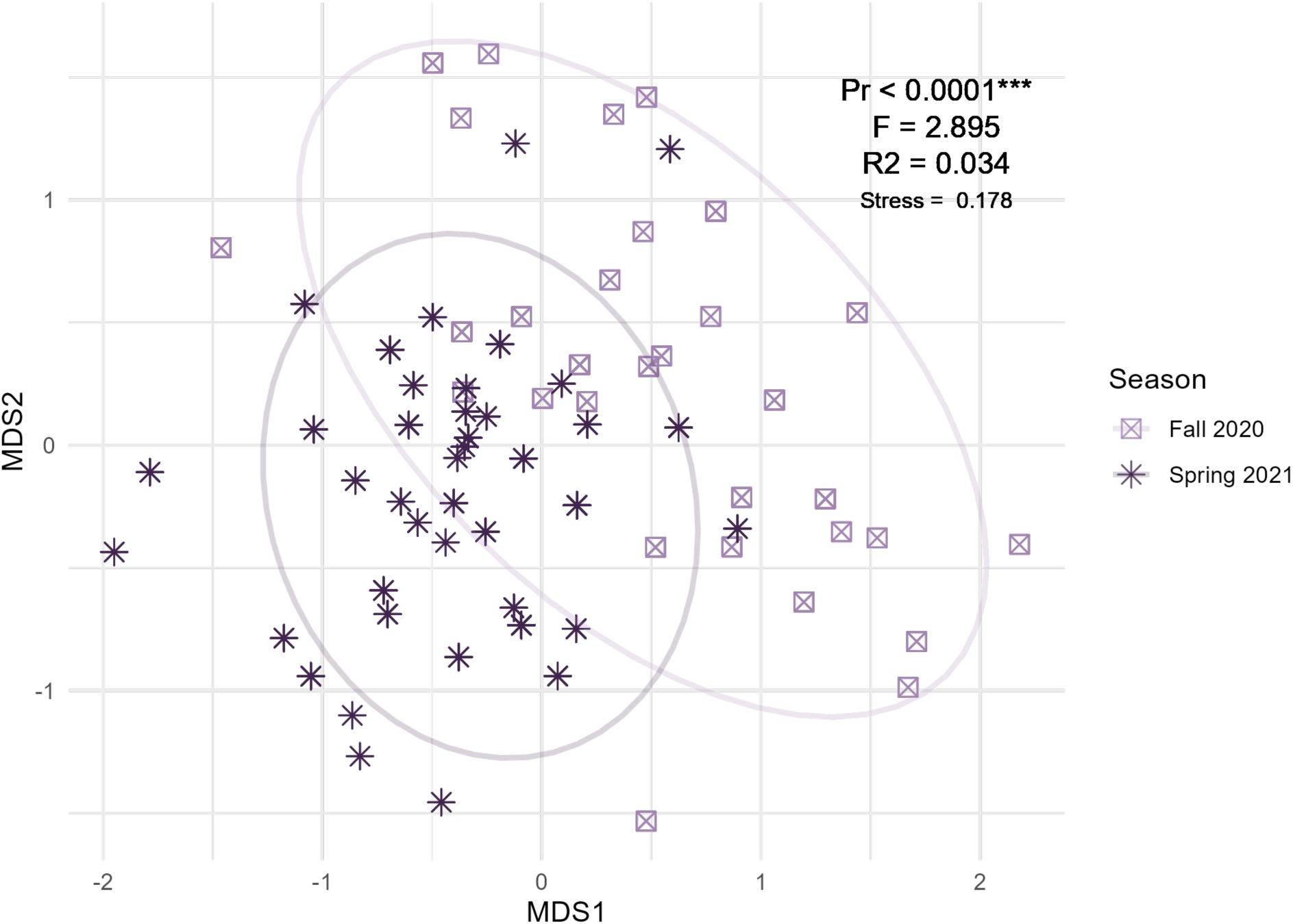
NMDS plot showing similarity of arthropod communities between sites, colored by season with Fall sites in light purple and Spring sites in dark purple. There was a significant difference by season.

**Figure 6.**
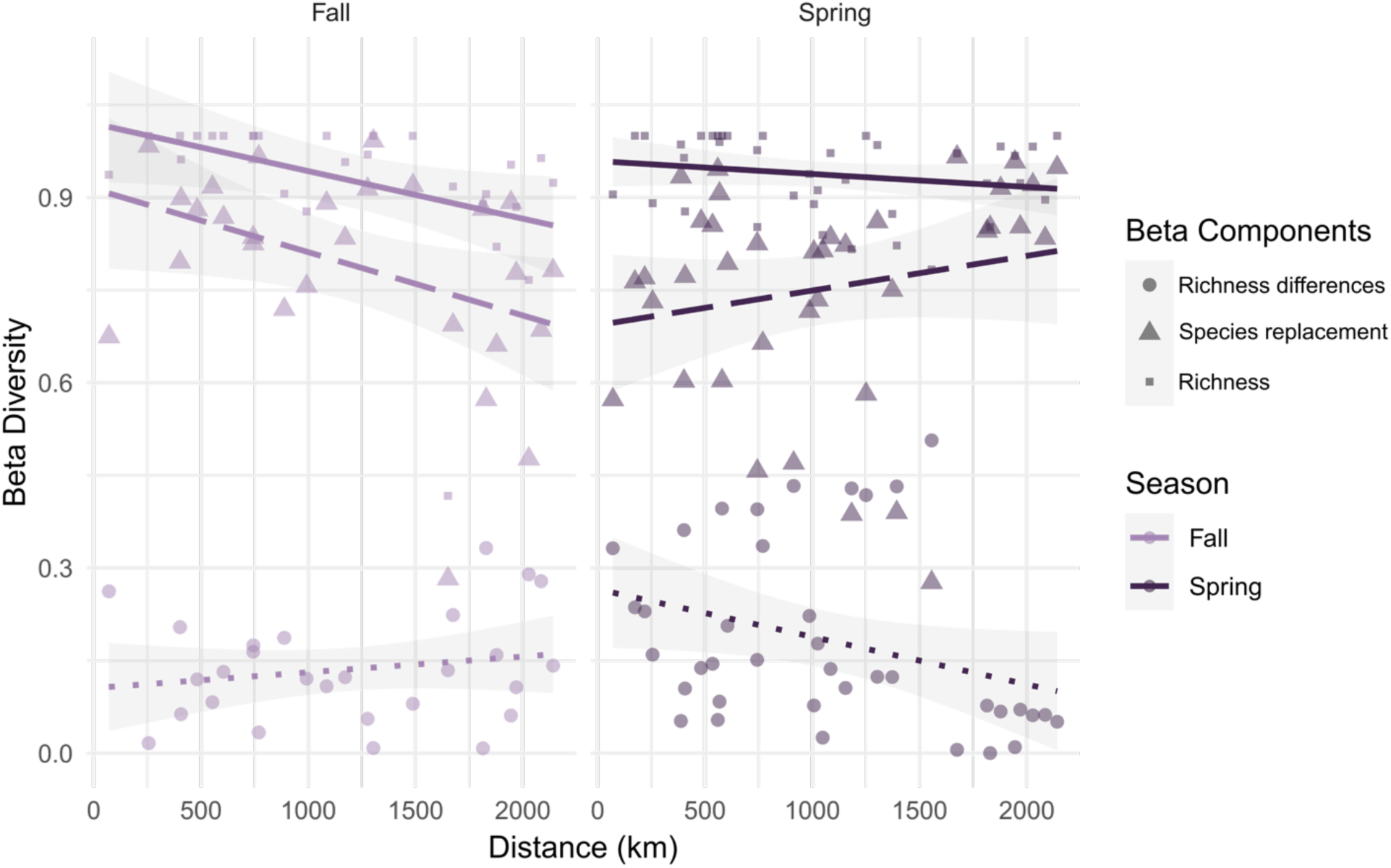
In the Fall (left), beta diversity values decreased with distance, largely due to decreases in the replacement component. This differed in the Spring (right), where the importance of replacement increased with distance.

### H3. Response by habitat type

Beta diversity between burned and unburned sites within habitat types did not vary, nor did they significantly change from Fall to Spring. In grassland, burn status was not explanatory in either season. However, there were significant dispersion differences in the Spring season (Pr = 0.006), which is also noticeable in the ordination plot (Figure 7a). Oak woodland sites show a similar pattern, without significant differences between groups in either year. Unlike grassland sites, no dispersion differences were detected in oak woodlands; sites show a similar spread in ordination space (Figure 7b). In scrub habitat, burn status was not explanatory in the fall but was a significant explanatory variable in the Spring (R2 = 0.103, F = 1.38, Pr = 0.029; Figure 7c). Points separate clearly in ordination space into two groups, especially noticeable in the Spring. There were no detectable dispersion differences in either season in scrub habitat. In scrub, the lowest median beta diversity, or the highest compositional similarity, was between burned habitats (median = 0.881) while the highest median beta diversity score was comparing burned and unburned sites (median = 0.972). This contrasted the other habitat types, with which the highest median beta diversity score was comparing between the burned sites (median = 0.833 grassland; median = 0.990 oak woodlands) and the lowest median beta diversity was comparing between unburned sites (median = 0.958 grassland; median = 0.918 oak woodlands) (Figure 8).

**Figure 7.**
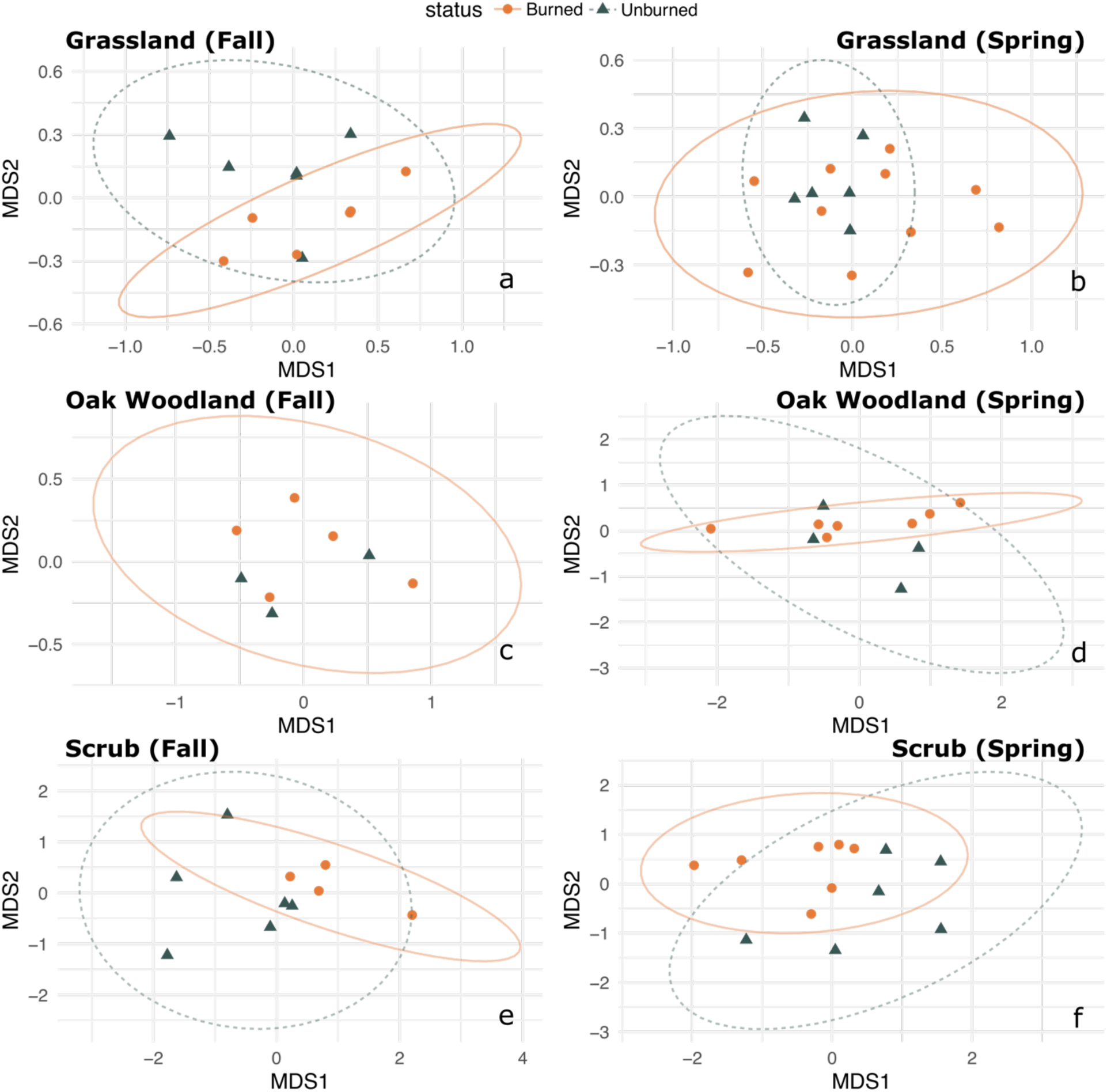
NMDS plots by habitat, with the top plots (a, b) showing grasslands, the middle plots (c, d) showing oak woodland and the bottom plots (e, f) showing scrub. Grassland sites became more similar in Spring (b), although burned sites showed significantly different dispersion. Oak woodlands were similar with burned and unburned clustering together in both season (c, d). There was a significant difference by burned and unburned status in scrub habitat for both seasons (e, f).

**Figure 8.**
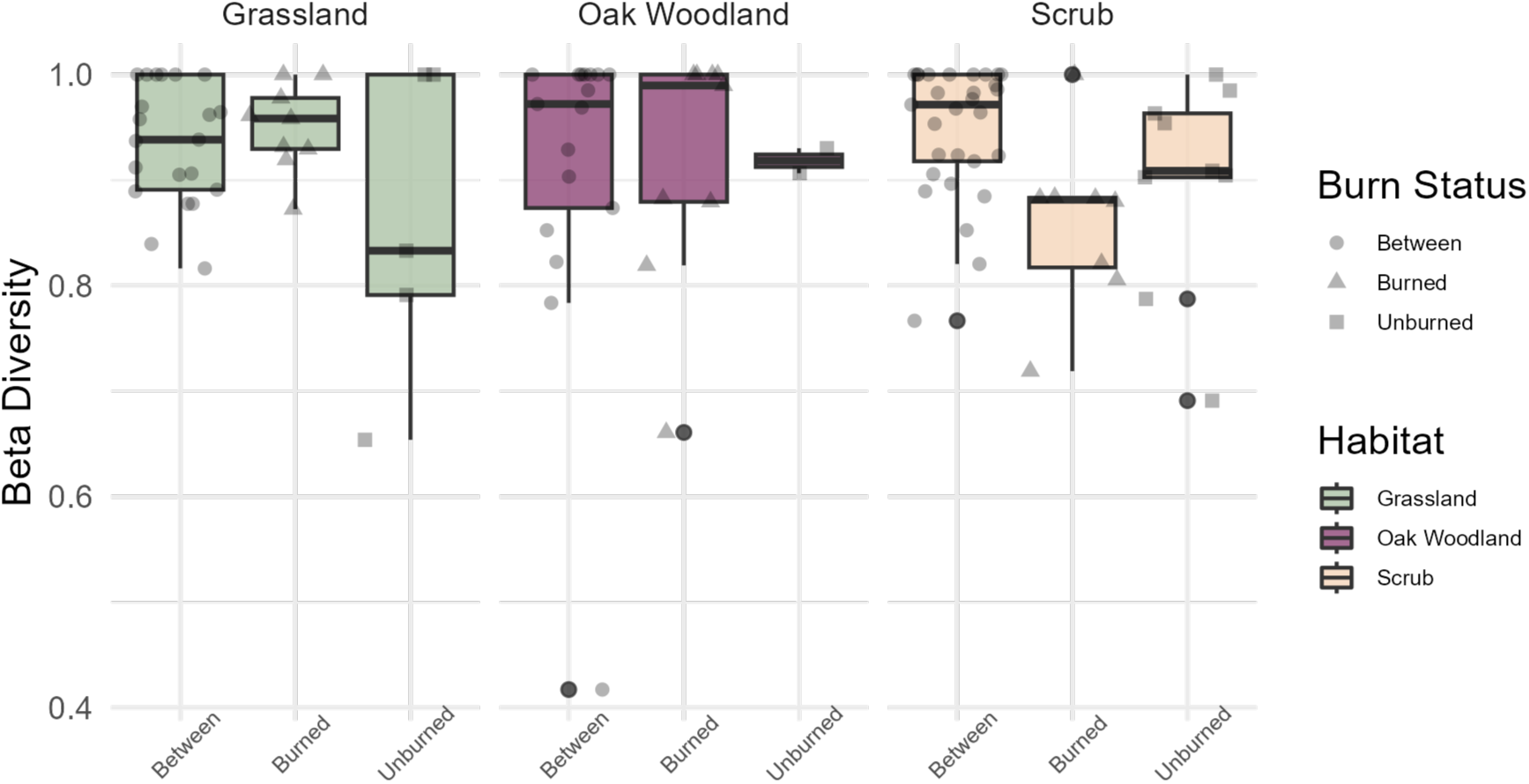
Beta diversity values split by comparisons between burned and unburned sites (Between), comparisons between sites only within burned habitat (Burned) and comparison between sites only in unburned habitat (Unburned). In scrub habitat, burned sites showed the lowest dissimilarity. The lowest dissimilarity score within grasslands and scrub habitats was within the comparison between unburned sites.

## Discussion

### H1. Alpha diversity differences between communities are explained largely by season rather than burn status

In this study, we utilized metabarcoding of whole arthropod communities collected from burned and unburned sites to assess the effect of large-scale wildfire on community composition across habitat types. Cyclic wildfires can foster biodiversity and many studies detect similar or higher species richness values in burned sites compared to unburned (Pausas and Parr 2018; Yekwayo et al. 2018). Our results using whole biotic communities align with this common finding, with no significant difference in alpha diversity values between burned and unburned sites (Figure 3). Our results show strong seasonal differences in alpha diversity (Figure 3; Figure 5). This supports our first hypothesis: alpha diversity values would not show differences by burn but would show differences by season. Certain arthropod species are pyrophilous, drawn to burned habitat by smoke or heat (Saint-Germain, Drapeau, and Buddle 2008). Immediate colonization during primary succession usually occurs by these fire-associated species followed by colonization by taxa that benefit indirectly from fire effects (Koltz et al. 2018; Boulanger and Sirois 2007). The emergence of plant species in the Spring, be it species commonly detected in the original ecosystem pre-burn or new species that capitalize on the burn, increases pollinator richness at all sites.

### H2. Communities showed strong compositional dissimilarity when comparing across reserves

While maintenance of biodiversity itself is important, it is essential to test patterns of species replacement and reordering if we are to understand how ecosystems change under novel disturbance. Burn status did not emerge as a significant grouping factor when comparing across all sites (Figure 4). Instead, we detected a highly significant difference in community composition based on reserve. Distance is then the most crucial factor in maintaining high diversity, with each reserve hosting a distinct pool of arthropods. This is true despite most taxa we collected being winged and presumably more capable of dispersal. Our second hypothesis predicted high connectivity across reserves within similar habitat types, but this finding counters our hypothesis. The high spatial turnover in diversity between reserves speaks to California’s classification as a biodiversity hotspot (Norman 2003). California is known for its incredibly diverse flora with around 6,900 native plants, most of which are endemic (Burge et al. 2016). Because insect diversity is often correlated with plant diversity (Novotny et al. 2006), we can expect that California has a high abundance of endemic insects with the distributions of many species’ closely associated with different plants. Protection of habitat more broadly is therefore essential in maintaining biodiversity.

### H3. The strength of community dissimilarities between burned and unburned sites were related to habitat type

To assess the effect of burn exclusively, we compared the compositions of sites within the same reserve, habitat type and season. Burn status was not a significant grouping factor itself when comparing across all sites. However, when reserve differences were modeled with an interaction term, burn status did emerge as an explanation of variation. Within reserves and the same habitat types, distance was significantly associated with beta diversity differences in the spring (Figure 6). Even on such a small scale (less than 3,000 meters), dispersal seems to be a limiting factor in recovery. The closer sites share more species-like groupings, with differences driven by richness, while the farther sites have beta diversity almost entirely driven by replacement. Interestingly, this trend is reversed in the fall. This may be explainable by the reduced alpha diversity in general, with taxa that remain active in the Fall being detected more broadly. The most abundant order in the Fall was Diptera (flies), which are less likely reliant on plant communities. Additionally, flies could potentially be more dispersive than Hymenoptera (ants, bees and wasps), which was the dominant order in the Spring and function more as the primary pollinators. The increased dissimilarity between sites in the spring could indicate that the successional trajectory is moving away from pre-fire conditions rather than towards. Long-lasting compositional differences are noted in other systems (Morris et al. 2019); these major changes in species composition can lead to an overall functionally distinct community (Podgaiski et al. 2013; Polchaninova et al. 2019). In places such as California with endemic taxa restricted to certain habitat types and demonstrating high spatial turnover, conversion of areas to novel ecosystems is a serious threat to maintenance of biodiversity.

The tight association between insects and flora makes consideration of habitat type crucial when measuring compositional response to fire. We found a significant interaction between habitat and reserve in explaining compositional differences between sites, emphasizing the importance of habitat type in biotic community structure. More dispersive and generalist insects may be favored following fire as they can capitalize on a novel plant community and colonize quickly (Koltz et al. 2018). Depending on dynamics of the wildfire, native vegetation can re-establish and flourish or instead be entirely missing from the plant community. Wildfires in forested areas can create a heterogeneous landscape with open areas that promote high flowering plant density and an associated increase in pollinators (Galbraith et al. 2019). Alternatively, frequent fires can cause invasion of non-native plants, particularly grasses, that can displace native species (Reilly et al. 2020; Hunter et al. 2006). Insects with particular specializations on endemic plant flora may not recover due to loss of host plants or competitive exclusion by opportunistic species that establish following fire.

The way a habitat will respond is driven both by the historical adaptation to fire and the particular dynamics of the wildfire. Grasslands are one habitat type that quickly flourishes after burn, regardless of severity. Within grasslands, non-native species of grasses are noted to become more prevalent following large-scale disturbance. Our results show that arthropod communities found in burned grassland sites resemble unburned sites by the following Spring (Figure 7b). This has been found in other groups, such as spiders, which show a similarly rapid recovery time-frame in grasslands following disturbance (Podgaiski et al. 2013). When comparing compositions within the burned sites, we see higher dissimilarities than observed between unburned sites. This may reflect the higher richness found in some of the burned grassland sites, and possibly reflects the benefits fire, even when unusually severe, can have on community diversity in certain habitat types.

Oak woodlands are another Californian habitat type reliant on regular wildfires in short intervals, which reduce density of conifer competitors and other fire intolerant species. Historical fire suppression caused major compositional changes to oak woodland habitat but prescribed burn strategies in more recent years have allowed these habitats to flourish once more. Oak woodlands are thought to be resilient even in the face of high severity burns, although this may impact the long-term development of the trees (Nemens et al. 2018). Our results in oak woodland sites showed no strong explanatory nature of burned and unburned in either fall or spring (Figure 7c,d). This aligns with the long history of fire in these habitat types, consisting of communities that are well-adapted and resilient to fire. Prescribed burn management over the last few decades has supported the naturally occurring biodiversity of oak woodlands. As in grasslands, the highest beta diversity scores in oak woodlands are when comparing between burned sites and when comparing between burned and unburned sites. Similarly, this may reflect the beneficial nature of fires in adapted systems.

While prescribed fire has been a regular part of management, application of standard fire management strategies in habitats with alternate fire regimes like scrubland will have consequential ecological effects (Calhoun et al. 2022). Fires are infrequent in scrubland, with intervals between 325 – 450 years for low sagebrush (Baker 2006). Despite this irregularity, scrublands were the most frequently burned habitat type from 2000 to 2020 in California^49^. Because early successional stages are more sensitive to perturbations, fire prior to robust recovery will alter the successional trajectory (Kröel-Dulay et al. 2015). Our results show that arthropod communities in burned scrub become more dissimilar than unburned sites in Spring (Figure 7f). Compared to both grasslands and oak woodlands, burned scrub sites show the lowest beta diversity scores between one another, potentially indicative of a homogenization process cause by novel disturbance regimes (Figure 8). This may be driven by the openness created by fire in scrub habitat that prompts establishment of non-native grasses and creates habitat that promotes establishment of species less common in scrub. Species that serve as bioindicators in scrub habitat, such as the greater sage-grouse (*Centrocercus urophasianus*), are strongly affected by large-scale wildfires, possibly because of increases in predator populations which benefit from the new vegetation (Anthony et al. 2022). Predator-prey densities also change, altering trophic networks (Holbrook et al. 2016). The recently established shorter fire intervals (less than five years) will likely prevent full recovery of scrubland and permanently alter ecosystem structure, shifting to more grassland-like habitat.

## Conclusions

Our study emphasizes the importance of studying the effect of wildfires across a broad spatial scale and diverse habitat types to better understand the nuanced community responses of ecosystems. As is the case with novel environmental changes, taxa must adapt, disperse or go extinct. For taxa dependent on fire, their adaptations are associated with aspects of the regime itself rather than general fire pressure, making new wildfire dynamics a novel pressure (Pausas and Keeley 2009). The limited distribution and extent of habitat such as scrublands provides few refugia for the endemic and endangered species located in scrublands threatened by wildfire. Empty niche space created by dispersing or locally extinct taxa will prompt establishment of non-native or opportunistic taxa and may permanently alter the successional trajectory of the habitat. Management methods which can assist in recovery following wildfires are essential to maintain such habitat types.

There are many future steps needed to make ecologically-informed management decisions. We took advantage of a natural experiment to build knowledge on whole community responses using a novel metabarcoding approach. Our study was limited by lack of taxonomic information available. More functional information is crucial to determine the cost of compositional differences for ecosystems. A larger reference collection of DNA barcodes verified at the species level would enable finer grained analysis of community dynamics including effect of fire on native and non-native species as well as restricting of the ecological network. Combining these data with attributes of the fire itself, associated with both historical and current fire regimes, will allow us to identify threatened ecosystems and move towards supporting recovery. With this information, we can more effectively assess community vulnerability and resilience in the face of severe climate events.

## Acknowledgements

We would like to thank Gage Dayton, Admin Director of UCSC Natural Reserves, and Kelly Zilliacus from the Conservation Action Lab at UCSC, for their assistance with the project and in coordinating collections. We would additionally like to thank Brandt Weary for his efforts in collecting samples. Our research would not have been possible without funding from UCOP and the California Heartbeat Initiative (CHI). We thank Becca Fenwick, the California Heartbeat Initiative program director, and Peggy Fiedler, former executive director of the UCNRS, for their assistance. Additional funds came from Margaret C. Walker Fund for Systematic Entomology as well as funds associated with the Schlinger Chair of Arachnology.

## Data availability

All relevant data and associated code are hosted on GitHub at https://github.com/rjcmarkelz/edna-fire. A Docker container exists to replicate processing of sequence data and can be found at https://hub.docker.com/r/codymarkelz/holmquist-etal-edna-fire-paper.

## References

Anthony, Christopher R., Lee J. Foster, Christian A. Hagen, and Katie M. Dugger. 2022. “Acute and Lagged Fitness Consequences for a Sagebrush Obligate in a Post Mega-Wildfire Landscape.” Ecology and Evolution 12 (1): e8488. https://doi.org/10.1002/ece3.8488.

Baker, William L. 2006. “Fire and Restoration of Sagebrush Ecosystems.” Wildlife Society Bulletin 34 (1): 177–85. https://doi.org/10.2193/0091-7648(2006)34[177:FAROSE]2.0.CO;2.

Barton, Andrew M. 2002. “Intense Wildfire in Southeastern Arizona: Transformation of a Madrean Oak–Pine Forest to Oak Woodland.” Forest Ecology and Management 165 (1): 205–12. https://doi.org/10.1016/S0378-1127(01)00618-1.

Beng, Kingsly Chuo, Kyle W. Tomlinson, Xian Hui Shen, Yann Surget-Groba, Alice C. Hughes, Richard T. Corlett, and J. W. Ferry Slik. 2016. “The Utility of DNA Metabarcoding for Studying the Response of Arthropod Diversity and Composition to Land-Use Change in the Tropics.” Scientific Reports 6 (1): 24965. https://doi.org/10.1038/srep24965.

Bieber, Blyssalyn V., Dhaval K. Vyas, Amanda M. Koltz, Laura A. Burkle, Kiaryce S. Bey, Claire Guzinski, Shannon M. Murphy, and Mayra C. Vidal. 2022. “Increasing Prevalence of Severe Fires Change the Structure of Arthropod Communities: Evidence from a Meta-Analysis.” Functional Ecology n/a (n/a). https://doi.org/10.1111/1365-2435.14197.

Boulanger, Yan, and Luc Sirois. 2007. “Postfire Succession of Saproxylic Arthropods, with Emphasis on Coleoptera, in the North Boreal Forest of Quebec.” Environmental Entomology 36 (1): 128–41. https://doi.org/10.1603/0046-225X-36.1.128.

Burge, Dylan O., James H. Thorne, Susan P. Harrison, Bart C. O’Brien, Jon P. Rebman, James R. Shevock, Edward R. Alverson, et al. 2016. “Plant Diversity and Endemism in the California Floristic Province.” Madroño 63 (2): 3–206. https://doi.org/10.3120/madr-63-02-3-206.1.

CAL Fire. 2022. “Top 20 Largest California Wildfires.”

Calhoun, Kendall L., Melissa Chapman, Carmen Tubbesing, Alex McInturff, Kaitlyn M. Gaynor, Amy Van Scoyoc, Christine E. Wilkinson, Phoebe Parker-Shames, David Kurz, and Justin Brashares. 2022. “Spatial Overlap of Wildfire and Biodiversity in California Highlights Gap in Non-Conifer Fire Research and Management.” Diversity and Distributions 28 (3): 529–41. https://doi.org/10.1111/ddi.13394.

Callahan, Benjamin J, Paul J McMurdie, Michael J Rosen, Andrew W Han, Amy Jo A Johnson, and Susan P Holmes. 2016. “DADA2: High-Resolution Sample Inference from Illumina Amplicon Data.” Nature Methods 13 (7): 581–83. https://doi.org/10.1038/nmeth.3869.

Carbone, Lucas M., Julia Tavella, Juli G. Pausas, and Ramiro Aguilar. 2019. “A Global Synthesis of Fire Effects on Pollinators.” Global Ecology and Biogeography 28 (10): 1487–98. https://doi.org/10.1111/geb.12939.

Cardoso, Pedro, François Rigal, and José C. Carvalho. 2015. “BAT – Biodiversity Assessment Tools, an R Package for the Measurement and Estimation of Alpha and Beta Taxon, Phylogenetic and Functional Diversity.” Edited by Steven Kembel. Methods in Ecology and Evolution 6 (2): 232–36. https://doi.org/10.1111/2041-210X.12310.

Connell, Joseph H. 1978. “Diversity in Tropical Rain Forests and Coral Reefs: High Diversity of Trees and Corals Is Maintained Only in a Nonequilibrium State.” Science 199 (4335): 1302–10. https://doi.org/10.1126/science.199.4335.1302.

Coop, Jonathan D, Sean A Parks, Camille S Stevens-Rumann, Shelley D Crausbay, Philip E Higuera, Matthew D Hurteau, Alan Tepley, et al. 2020. “Wildfire-Driven Forest Conversion in Western North American Landscapes.” BioScience 70(8): 659–73. https://doi.org/10.1093/biosci/biaa061.

Davis, Nicole M., Diana M. Proctor, Susan P. Holmes, David A. Relman, and Benjamin J. Callahan. 2018. “Simple Statistical Identification and Removal of Contaminant Sequences in Marker-Gene and Metagenomics Data.” Microbiome 6 (1): 226. https://doi.org/10.1186/s40168-018-0605-2.

Dürrbaum, Ellen, Felix Fornoff, Christoph Scherber, Eero J. Vesterinen, and Bernhard Eitzinger. n.d. “Metabarcoding of Trap Nests Reveals Differential Impact of Urbanization on Cavity-Nesting Bee and Wasp Communities.” Molecular Ecology n/a (n/a). Accessed July 21, 2023. https://doi.org/10.1111/mec.16818.

Eds, Core Writing Team. 2007. “Contribution of Working Groups I, II and III to the Fourth Assessment Report of the Intergovernmental Panel on Climate Change.” IPCC 2007 : Climate Change 2007 : Synthesis Report 104. https://cir.nii.ac.jp/crid/1571417124957557504.

Elbrecht, Vasco, Thomas W. A. Braukmann, Natalia V. Ivanova, Sean W. J. Prosser, Mehrdad Hajibabaei, Michael Wright, Evgeny V. Zakharov, Paul D. N. Hebert, and Dirk Steinke. 2019. “Validation of COI Metabarcoding Primers for Terrestrial Arthropods.” PeerJ 7 (October): e7745. https://doi.org/10.7717/peerj.7745.

Frøslev, Tobias Guldberg, Rasmus Kjøller, Hans Henrik Bruun, Rasmus Ejrnæs, Ane Kirstine Brunbjerg, Carlotta Pietroni, and Anders Johannes Hansen. 2017. “Algorithm for Post-Clustering Curation of DNA Amplicon Data Yields Reliable Biodiversity Estimates.” Nature Communications 8 (1): 1188. https://doi.org/10.1038/s41467-017-01312-x.

Galbraith, Sara M., James H. Cane, Andrew R. Moldenke, and James W. Rivers. 2019. “Wild Bee Diversity Increases with Local Fire Severity in a Fire-Prone Landscape.” Ecosphere 10 (4): e02668. https://doi.org/10.1002/ecs2.2668.

Holbrook, Joseph D., Robert S. Arkle, Janet L. Rachlow, Kerri T. Vierling, David S. Pilliod, and Michelle M. Wiest. 2016. “Occupancy and Abundance of Predator and Prey: Implications of the Fire-Cheatgrass Cycle in Sagebrush Ecosystems.” Ecosphere 7 (6): e01307. https://doi.org/10.1002/ecs2.1307.

Holmquist, Anna J., Seira A. Adams, and Rosemary G. Gillespie. 2023. “Invasion by an Ecosystem Engineer Changes Biotic Interactions between Native and Non-Native Taxa.” Ecology and Evolution 13 (2): e9820. https://doi.org/10.1002/ece3.9820.

Hunter, Molly E., Philip N. Omi, Erik J. Martinson, Geneva W. Chong, Molly E. Hunter, Philip N. Omi, Erik J. Martinson, and Geneva W. Chong. 2006. “Establishment of Non-Native Plant Species after Wildfires: Effects of Fuel Treatments, Abiotic and Biotic Factors, and Post-Fire Grass Seeding Treatments.” International Journal of Wildland Fire 15 (2): 271–81. https://doi.org/10.1071/WF05074.

Jager, Henriette I., Jonathan W. Long, Rachel L. Malison, Brendan P. Murphy, Ashley Rust, Luiz G. M. Silva, Rahel Sollmann, et al. 2021. “Resilience of Terrestrial and Aquatic Fauna to Historical and Future Wildfire Regimes in Western North America.” Ecology and Evolution 11 (18): 12259–84. https://doi.org/10.1002/ece3.8026.

Johnstone, Jill F, Craig D Allen, Jerry F Franklin, Lee E Frelich, Brian J Harvey, Philip E Higuera, Michelle C Mack, et al. 2016. “Changing Disturbance Regimes, Ecological Memory, and Forest Resilience.” Frontiers in Ecology and the Environment 14 (7): 369–78. https://doi.org/10.1002/fee.1311.

Koltz, Amanda M, Laura A Burkle, Yamina Pressler, Jane E Dell, Mayra C Vidal, Lora A Richards, and Shannon M Murphy. 2018. “Global Change and the Importance of Fire for the Ecology and Evolution of Insects.” *Current Opinion in Insect Science*, Global change biology * Molecular physiology, 29 (October): 110–16. https://doi.org/10.1016/j.cois.2018.07.015.

Kröel-Dulay, György, Johannes Ransijn, Inger Kappel Schmidt, Claus Beier, Paolo De Angelis, Giovanbattista de Dato, Jeffrey S. Dukes, et al. 2015. “Increased Sensitivity to Climate Change in Disturbed Ecosystems.” Nature Communications 6 (1): 6682. https://doi.org/10.1038/ncomms7682.

Lecina-Diaz, Judit, Jordi Martínez-Vilalta, Albert Alvarez, Jordi Vayreda, and Javier Retana. 2021. “Assessing the Risk of Losing Forest Ecosystem Services Due to Wildfires.” Ecosystems 24 (7): 1687–1701. https://doi.org/10.1007/s10021-021-00611-1.

Leray, Matthieu, Joy Y. Yang, Christopher P. Meyer, Suzanne C. Mills, Natalia Agudelo, Vincent Ranwez, Joel T. Boehm, and Ryuji J. Machida. 2013. “A New Versatile Primer Set Targeting a Short Fragment of the Mitochondrial COI Region for Metabarcoding Metazoan Diversity: Application for Characterizing Coral Reef Fish Gut Contents.” Frontiers in Zoology 10 (1): 34. https://doi.org/10.1186/1742-9994-10-34.

Martin, Marcel. 2011. “Cutadapt Removes Adapter Sequences from High-Throughput Sequencing Reads.” EMBnet. Journal 17 (1): 10–12. https://doi.org/10.14806/ej.17.1.200.

Moritz, Max A., Marc-André Parisien, Enric Batllori, Meg A. Krawchuk, Jeff Van Dorn, David J. Ganz, and Katharine Hayhoe. 2012. “Climate Change and Disruptions to Global Fire Activity.” Ecosphere 3 (6): art49. https://doi.org/10.1890/ES11-00345.1.

Morris, Jesse L., R. Justin DeRose, Thomas Brussel, Simon Brewer, Andrea Brunelle, and James N. Long. 2019. “Stable or Seral? Fire-Driven Alternative States in Aspen Forests of Western North America.” Biology Letters 15 (6): 20190011. https://doi.org/10.1098/rsbl.2019.0011.

Nemens, Deborah G., J. Morgan Varner, Kathryn R. Kidd, and Brian Wing. 2018. “Do Repeated Wildfires Promote Restoration of Oak Woodlands in Mixed-Conifer Landscapes?” Forest Ecology and Management 427 (November): 143–51. https://doi.org/10.1016/j.foreco.2018.05.023.

Norman, Myers. 2003. “Biodiversity Hotspots Revisited.” BioScience 53 (10): 916–17. https://doi.org/10.1641/0006-3568(2003)053[0916:BHR]2.0.CO;2.

Novotny, Vojtech, Pavel Drozd, Scott E. Miller, Miroslav Kulfan, Milan Janda, Yves Basset, and George D. Weiblen. 2006. “Why Are There So Many Species of Herbivorous Insects in Tropical Rainforests?” Science 313 (5790): 1115–18. https://doi.org/10.1126/science.1129237.

Oksanen, Jari, Roeland Kindt, Pierre Legendre, Bob Hara, Gavin Simpson, Peter Solymos, M Henry, Hank Stevens, Helene Maintainer, and jari Oksanen@oulu. 2009. “The Vegan Package,” January.

Parks, S. A., and J. T. Abatzoglou. 2020. “Warmer and Drier Fire Seasons Contribute to Increases in Area Burned at High Severity in Western US Forests From 1985 to 2017.” Geophysical Research Letters 47 (22): e2020GL089858. https://doi.org/10.1029/2020GL089858.

Pausas, Juli G., and Jon E Keeley. 2009. “Burning Story: The Role of Fire in the History of Life.” BioScience 59. https://academic.oup.com/bioscience/article/59/7/593/334816.

Pausas, Juli G, and Jon E Keeley. 2019. “Wildfires as an Ecosystem Service.” Frontiers in Ecology and the Environment 17 (5): 289–95. https://doi.org/10.1002/fee.2044.

Pausas, Juli G., and Catherine L. Parr. 2018. “Towards an Understanding of the Evolutionary Role of Fire in Animals.” Evolutionary Ecology 32 (2): 113–25. https://doi.org/10.1007/s10682-018-9927-6.

Podgaiski, Luciana R., Fernando Joner, Sandra Lavorel, Marco Moretti, Sebastien Ibanez, Milton De S. Mendonça, and Valério D. Pillar. 2013. “Spider Trait Assembly Patterns and Resilience under Fire-Induced Vegetation Change in South Brazilian Grasslands.” Edited by Nathan G. Swenson. PLoS ONE 8 (3): e60207. https://doi.org/10.1371/journal.pone.0060207.

Polchaninova, Nina, Galina Savchenko, Vladimir Ronkin, Aleksandr Drogvalenko, and Alexandr Putchkov. 2019. “Summer Fire in Steppe Habitats: Long-Term Effects on Vegetation and Autumnal Assemblages of Cursorial Arthropods.” Hacquetia 18 (2): 213–31. https://doi.org/10.2478/hacq-2019-0006.

Ponisio, Lauren C., Kate Wilkin, Leithen K. M’Gonigle, Kelly Kulhanek, Lindsay Cook, Robbin Thorp, Terry Griswold, and Claire Kremen. 2016. “Pyrodiversity Begets Plant–Pollinator Community Diversity.” Global Change Biology 22 (5): 1794–1808. https://doi.org/10.1111/gcb.13236.

Reilly, Matthew J., Millen G. McCord, Stefani M. Brandt, Kevin P. Linowksi, Ramona J. Butz, and Erik S. Jules. 2020. “Repeated, High-Severity Wildfire Catalyzes Invasion of Non-Native Plant Species in Forests of the Klamath Mountains, Northern California, USA.” Biological Invasions 22 (6): 1821–28. https://doi.org/10.1007/s10530-020-02227-3.

Roxburgh, S.H., K Shea, and Wilson, J.B. 2004. “The Intermediate Disturbance Hypothesis: Patch Dynamics and Mechanisms of Species Coexistence.” Ecology 85: 359–71.

Saint-Germain, Michel, Pierre Drapeau, and Christopher M. Buddle. 2008. “Persistence of Pyrophilous Insects in Fire-Driven Boreal Forests: Population Dynamics in Burned and Unburned Habitats.” Diversity and Distributions 14 (4): 713–20. https://doi.org/10.1111/j.1472-4642.2007.00452.x.

Schoennagel, Tania, Jennifer K. Balch, Hannah Brenkert-Smith, Philip E. Dennison, Brian J. Harvey, Meg A. Krawchuk, Nathan Mietkiewicz, et al. 2017. “Adapt to More Wildfire in Western North American Forests as Climate Changes.” Proceedings of the National Academy of Sciences 114 (18): 4582–90. https://doi.org/10.1073/pnas.1617464114.

Seidl, Rupert, Dominik Thom, Markus Kautz, Dario Martin-Benito, Mikko Peltoniemi, Giorgio Vacchiano, Jan Wild, et al. 2017. “Forest Disturbances under Climate Change.” Nature Climate Change 7 (6): 395–402. https://doi.org/10.1038/nclimate3303.

Taillie, Paul J., Ryan D. Burnett, Lance Jay Roberts, Brent R. Campos, M. Nils Peterson, and Christopher E. Moorman. 2018. “Interacting and Non-Linear Avian Responses to Mixed-Severity Wildfire and Time since Fire.” Ecosphere 9 (6): e02291. https://doi.org/10.1002/ecs2.2291.

Wilkinson, S. 2018. “Kmer: An R Package for Fast Alignment-Free Clustering of Biological.” Winter, David J. 2017. “Rentrez: An R Package for the NCBI EUtils API.” e3179v2. PeerJ Inc. https://doi.org/10.7287/peerj.preprints.3179v2.

Yekwayo, Inam, James S. Pryke, René Gaigher, and Michael J. Samways. 2018. “Only Multi-Taxon Studies Show the Full Range of Arthropod Responses to Fire.” PLOS ONE 13 (4): e0195414. https://doi.org/10.1371/journal.pone.0195414.

Yu, Douglas W., Yinqiu Ji, Brent C. Emerson, Xiaoyang Wang, Chengxi Ye, Chunyan Yang, and Zhaoli Ding. 2012. “Biodiversity Soup: Metabarcoding of Arthropods for Rapid Biodiversity Assessment and Biomonitoring.” Methods in Ecology and Evolution 3 (4): 613–23. https://doi.org/10.1111/j.2041-210X.2012.00198.x.

