## Supplemental Materials for "The importance of habitat type and historical fire regimes in arthropod community response following large-scale wildfires"

### Supplementary Materials

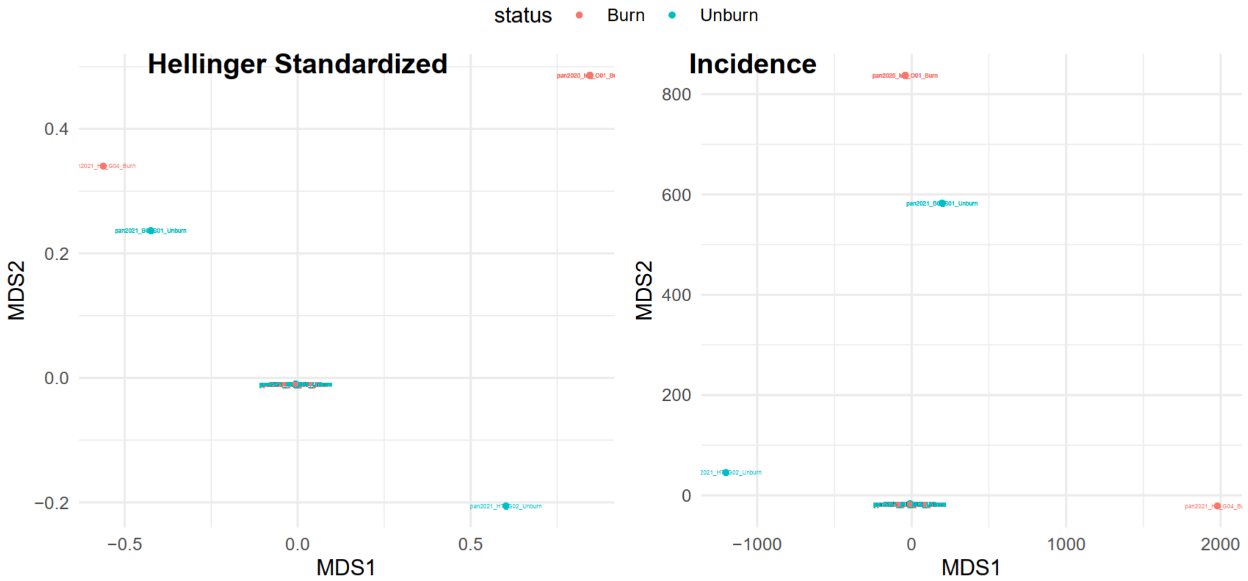


**Supplementary Materials, Figure 1.** Outlier samples with few OTUs that matched no other samples, causing convergence issues in NMDS.

| Predictor | SumOfSqs | R2 | F | Pr(>F) |
|---|---|---|---|---|
| year | 1.3127523 | 0.03300788 | 3.228144 | 0.000999001 |
| status | 0.5209520 | 0.01309883 | 1.281055 | 0.025974026 |
| habitat | 1.2889699 | 0.03240989 | 1.584830 | 0.000999001 |
| reserve | 3.1862222 | 0.08011446 | 1.958782 | 0.000999001 |
| year:status | 0.4926112 | 0.01238623 | 1.211363 | 0.085914086 |
| year:habitat | 1.0459904 | 0.02630041 | 1.286079 | 0.004995005 |
| status:habitat | 0.9230291 | 0.02320867 | 1.134894 | 0.102897103 |
| year:reserve | 2.9792902 | 0.07491136 | 1.831567 | 0.000999001 |
| status:reserve | 2.0334191 | 0.05112835 | 1.250077 | 0.005994006 |
| habitat:reserve | 1.9683661 | 0.04949265 | 1.210085 | 0.004995005 |
| year:status:habitat | 0.8712574 | 0.02190692 | 1.071239 | 0.230769231 |
| year:status:reserve | 1.8830717 | 0.04734801 | 1.157649 | 0.015984016 |
| year:habitat:reserve | 1.7853202 | 0.04489014 | 1.097555 | 0.074925075 |
| status:habitat:reserve | 1.1757126 | 0.02956215 | 0.963718 | 0.672327672 |
| year:status:habitat:reserve | 0.8175907 | 0.02055752 | 1.005254 | 0.451548452 |
| Residual | 17.4863199 | 0.43967652 | NA | NA |
| Total | 39.7708750 | 1.00000000 | NA | NA |

**Supplementary materials, Table 1.** Results of PERMANOVA using Jaccard dissimilarity (incidence data) and inclusion of outliers.

| Predictor | SumOfSqs | R2 | F | Pr(>F) |
|---|---|---|---|---|
| year | 2.1660164 | 0.02687548 | 2.5602735 | 0.000999001 |
| status | 0.9836265 | 0.01220463 | 1.1626657 | 0.108891109 |
| habitat | 2.4344191 | 0.03020576 | 1.4387653 | 0.000999001 |
| reserve | 6.3635555 | 0.07895767 | 1.8804616 | 0.000999001 |
| year:status | 0.9872088 | 0.01224908 | 1.1669000 | 0.081918082 |
| year:habitat | 2.2010716 | 0.02731044 | 1.3008547 | 0.002997003 |
| status:habitat | 1.8856528 | 0.02339679 | 1.1144391 | 0.100899101 |
| year:reserve | 5.6111939 | 0.06962252 | 1.6581351 | 0.000999001 |
| status:reserve | 4.0178322 | 0.04985242 | 1.1872889 | 0.008991009 |
| habitat:reserve | 3.9943578 | 0.04956116 | 1.1803521 | 0.009990010 |
| year:status:habitat | 1.9522288 | 0.02422285 | 1.1537862 | 0.043956044 |
| year:status:reserve | 3.6665315 | 0.04549356 | 1.0834779 | 0.091908092 |
| year:habitat:reserve | 3.7032873 | 0.04594962 | 1.0943394 | 0.069930070 |
| status:habitat:reserve | 2.5350106 | 0.03145388 | 0.9988106 | 0.489510490 |
| year:status:habitat:reserve | 1.7141061 | 0.02126827 | 1.0130534 | 0.414585415 |
| Residual | 36.3784208 | 0.45137586 | NA | NA |
| Total | 80.5945198 | 1.00000000 | NA | NA |

**Supplementary materials, Table 2.** Results of PERMANOVA using Euclidean distance matrix (Hellinger) and inclusion of outliers.
